# Extended Phenotype Influences Collective Behaviour and Survival in a Social Spider

**DOI:** 10.1101/2024.09.12.612246

**Authors:** Bharat Parthasarathy, Naeem Yusuf Shaikh, V Sai Abhinay, C Varun Sai, M V Sai Krishna, Krishna Kiran Vamsi Dasu

## Abstract

Increasing human interference has been shown to not only destroy habitats, but also alter the architecture of animal-built extended phenotypes. However, the impact of such architectural changes on the behaviour and survival of organisms remains poorly understood. To address this knowledge gap, we examined the impact of habitat modification using Indian social spider, *Stegodyphus sarasinorum*, as a model organism. *S. sarasinorum* colonies typically construct a three-dimensional (3D) capture web. Due to increasing habitat modification by humans, these spiders are now constrained to build two-dimensional (2D) capture webs adapting to man-made structures like fences. We investigated how these differing web architectures influence the collective behaviours and survival of *S. sarasinorum*. Our findings reveal that spiders with 2D capture webs emerged from their nests sooner, attacked prey faster, and had higher number of attacking spiders compared to those with 3D webs, suggesting 2D webs may be more efficient for hunting. However, despite their hunting advantages, spiders in 2D webs more frequently attacked the dangerous body parts of honeybees and were susceptible to honeybee stings. These results suggest that human-induced architectural modifications of the extended phenotype can have both benefits and costs for the organisms that built it. The survival benefits conferred by 3D capture webs against risky prey may have played a significant role in the evolutionary selection of this web architecture in *S. sarasinorum*.

## Introduction

Animals across diverse taxa construct impressive architectural structures (extended phenotypes) such as traps, pits, nests, and dams that have long fascinated biologists (Dawkins 1982). These animal-engineered extended phenotypes serve crucial functions, including providing protection, stabilizing temperature, facilitating aeration, trapping food, and potentially altering environmental selection pressures on the animals and their offspring (Clark et al. 2020; Laland et al. 2016; Gupta et al. 2017).

Animals have demonstrated the ability to modify their extended phenotypes in response to various intrinsic and extrinsic conditions (Farji-Brener and Amador-Vargas 2020; Ellandula et al. 2021). Interestingly, the components of these extended phenotypes can also underlie trade-offs. For example, the web-building behaviour of the social spider *Stegodyphus sarasinorum* reflects a classic trade-off between foraging and sheltering. Depending on its hunger, the spider adjusts its investment in protective nests versus cribellate prey capture webs (Parthasarathy et al. 2022). These findings demonstrate an interplay between an animal’s behaviour and the architectural structure and design of its extended phenotype.

While it is well-established that the environment can influence animal behaviours and fitness (Real 1994; Wong & Candolin 2015; Bernhardt et al 2020; Wilson et al. 2020), specifically whether the architectural design of extended phenotypes can, in-turn, influence an animal’s behaviour, fitness, and survival remains unclear. For example, a nest-building animal within a population which is more vulnerable to predators might build more defensive nests (Burtka and Grindstaff 2013; Trnka et al. 2013). However, it is unclear whether a less defensive nest architecture can in-turn prompt an individual to behave more aggressively against predators. Empirical evidence demonstrating that individual behaviours can in-turn be modulated by extended phenotypes are scarce: Pinter-Wollman (2015) found a positive relationship between nest chamber connectivity and foraging speed in harvester ants, while Montiglio and DiRienzo (2016) showed that web architecture contributed to the within-individual variation in foraging aggression in black widow spiders. Further research is needed to fully understand the feedback between extended phenotypes and behaviour, and how this dynamic affects survival and fitness.

The architectural design of a spider’s capture web can significantly influence its prey capture success (Rypstra 1982; Zschokke et al. 2006), survival (Blackledge et al. 2003), and the habitat preference (Haberkern et al. 2020). Notably, natural populations of *S. sarasinorum* exhibit both two-dimensional (2D) and three-dimensional (3D) web forms, providing the opportunity to investigate the collective behaviours and survival of spiders on these two web architectures. A previous research on the congeneric *Stegodyphus dumicola* found that individuals living in 2D nests hunted prey more effectively compared to individuals living in 3D nests (Kamath et al. 2019). 3D capture webs with sheets positioned along varying planes, may be less efficient at transmitting prey vibration cues to the spiders’ nests. Conversely, 2D webs with sheets along a single plane, can be less effective at ensnaring dangerous prey, potentially posing greater risks to the spiders or necessitating different hunting strategies.

In this study, we investigated how the architecture of extended phenotypes can shape the collective behaviours and survival of animals, using the Indian social spider *Stegodyphus sarasinorum* as a model organism. In the natural habitat, *S. sarasinorum* colonies inhabit plants and trees in the arid and semi-arid habitats of the Indian subcontinent. These spiders construct a tubular nest where they reside, as well as a cribellate capture web extending from the nest to trap prey. The capture webs consist of multiple flat web sheets that extend in all planes, forming a three-dimensional (3D) web architecture (Avilés 1997). Sociality has evolved multiple times in spiders (Agnarsson et al. 2006; Johannesen et al. 2007), and all social species except *Aebutina binotata* build 3D capture webs (Avilés 1993). This suggests that 3D web architecture is an important yet unexplored factor in the social evolution of spiders. However, with increasing human encroachment and habitat modification, *S. sarasinorum* colonies now build single, flat capture webs on man-made structures like fences and poles. The spatial constraints of these artificially-built structures prevent the webs from expanding in all planes, resulting in a two-dimensional (2D) architecture rather than the natural 3D form.

We investigated how web architecture affects the hunting behaviour of *S. sarasinorum* by setting up experimental colonies in the lab which either built 2D or 3D capture webs. We then analyzed how the web architecture impacted spider survival and the collective hunting strategies used by the spiders.

## Materials and Methods

### Study Organism

*Stegodyphus sarasinorum* (Karsch) is the sole social spider species found in the Indian subcontinent (Lubin & Bilde 2007). Adult colonies typically consist of 30-50 individuals, which work together to construct a web for capturing prey as well as a nest where the spiders live. *S. sarasinorum* capture webs are cobwebs with a typical three-dimensional structure, as opposed to two-dimensional orb webs (Avilés 1997). However, due to space and structural limitations, *S. sarasinorum* colonies has demonstrated the ability to construct two-dimensional capture webs and a more homogeneous nest on man-made structures like fences and poles due to space and/or structural limitations.

### Spider Collection and Construction of Experimental Colonies

We performed two different experiments at different periods using two different populations of *S. sarasinorum*. 1. We performed the first experiment (Experiment 1) in January and February 2020, at the Indian Institute of Science (IISC), Bangalore, India. We collected 20 colonies consisting of subadult *S. sarasinorum* throughout a 30-kilometer stretch near Kuppam, Andhra Pradesh, Southern India. We performed the second experiment (Experiment 2) in May and June 2024 at the Sri Sathya Sai Institute of Higher Learning (SSSIHL) in Puttaparthi, Andhra Pradesh, India. We collected 20 colonies consisting of subadult *S. sarasinorum* along a 45-kilometer length around Puttaparthi, Andhra Pradesh. The two populations were separated by an aerial distance of approximately 170 km.

We placed the collected colonies inside well-ventilated plastic containers (30 × 20 × 10 cm) and transported them to the respective laboratories. We randomly selected 40 spiders from each colony and uniquely marked them with non-toxic water colors (Camlin, Kokuyo Co. Ltd) for identification. We then transferred these spiders to either an aluminum frame (20 × 20 cm) or a wooden cage (20 × 20 × 20 cm) with mesh enclosures to simulate two- and three-dimensional environments (**Figure 1**) with a set of 10 colonies in each group. For the first experiment, the frames were made of wood instead of aluminum. Two colonies belonging to 2D frames in the first experiment perished for unknown reasons just before the trials began, so we were left with 18 colonies for the experiments. In the case of 2D frames, we covered the top corners of the frames with black chart paper to simulate a nest-like environment, as *S. sarasinorum* tend to settle and build nests in the top corners. Due to minimal depth of the frames, the spider colonies were restrained and could only construct 2D capture webs. To prevent spiders from escaping, we hung the frames 7 feet above the floor using a plastic rope which was applied with petroleum jelly. In contrast, the spiders in the wooden cages could construct 3D capture webs with the help of wooden sticks placed inside the cages. We allowed 7 days for the colonies to settle and build webs, during which we generously fed them honeybees (*Apis cerana*) and sprayed them with water daily.

**Figure 1:**
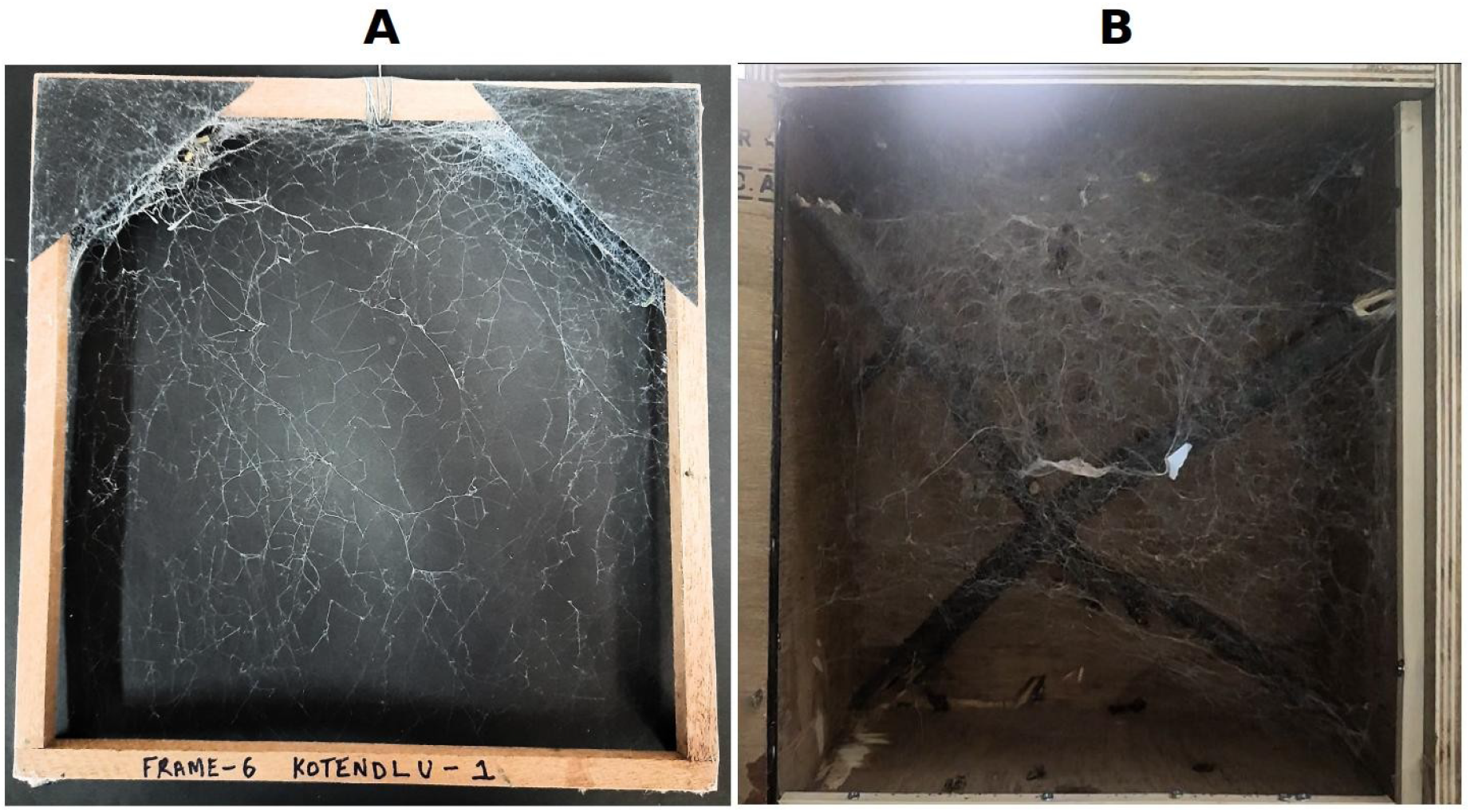
2D and 3D web architecture of *S. sarasinorum* in the laboratory. Shown here are a 2D colony on a frame (A) against a black background and a 3D colony inside a wooden box (B). The top corners of the frame were covered with chart paper to simulate a nest-like environment. The 2D colony constructs a flat sheet-like web constrained to a single plane by the frame. In contrast, the 3D colony inside the box is able to build sheet webs that extend across multiple planes.

### Prey Capture and Simulated Prey Capture Trials

#### Experiment 1

We used living, forager honeybees as prey in this experiment. We captured foragers from a beehive maintained in IISC campus and we gently placed them at the center of the spider’s web for both 2D and 3D experimental set-up. We noted the specific body parts of the bee that the spiders attacked, both before and for 1 minute after the bee was subdued. We categorized the attacks on the legs, wings and antenna of the honeybee as “attacks on the appendages” whereas attacks on the head, thorax and abdomen of the bee were categorized as “attacks on the body”. We also noted the number of spiders that succumbed to the bee stings while hunting. After each trial, we removed the dead bee to prevent the spiders from feeding on it. We performed 15 such trials over the course of 8 days. Subsequently, we reversed the web architecture of each colony by transferring the spiders from 2D frames and 3D cages and vice versa. We allowed the colonies another 7 days to settle and build capture webs, as described above. We then performed another set of 15 trials over 8 days.

#### Experiment 2

We used an electric handheld vibrator set to a specific frequency to simulate a struggling prey on the web. As mentioned previously, we placed the vibrator at the center of the web to simulate the presence of prey. We recorded the time (in *s*) it took for the spiders to emerge from the nest, followed by the time (in *s*) required to attack the vibrator string. We also noted the number of spiders that participated in the attack. We abandoned the trial if no response was noted within 2 minutes. We conducted 12 trials over 4 days. Subsequently, we reversed the web architecture as described in experiment 1, following which we performed another set of 12 trials over 4 days. Some colonies did not respond to the vibration for 12 trials (range for total trials responded: 4 – 12 trials; see **Supplementary Table 1** for colony-wise breakup). One colony, after reversal, perished for unknown reasons, so we could perform only 4 trials for this particular colony.

### Statistical Analyses

We conducted all analyses using the R programming language (R core team 2021). Abandoned trials in Experiment 2 were not included in the analyses. Since we used a before-after experimental design, we first had to rule out potential interaction effects of sequence (2D to 3D or 3D to 2D) and web architecture (2D or 3D webs). We built a generalized linear mixed effects model for this purpose using the lme4 package (Bates et al. 2015) in R. The response variables were latency for emergence, latency to attack, or number of attacking spiders. Sequence and web architecture were categorical fixed effects, while colony identity was a random effect. Finding no significant interaction between sequence and web architecture, we concluded that the order of experiments did not influence our results on latencies to emerge and attack or the number of attackers. Next, we built separate generalized linear mixed models. The continuous response variables (latency to emerge, latency to attack) were log-transformed to improve normality of residuals. For the model involving number of attacking spiders, we employed a Poisson regression mixed model. The fixed effects were web architecture, whether the experiment was reversed, and trial number, while colony identity was the random effect. We tested model assumptions regarding normality of residuals and inspected model fit by checking residuals plotted against observed values. We used Tukey’s post-hoc test to obtain pairwise comparisons of latencies to emerge and attack, as well as number of attacking spiders, between 2D and 3D colonies. Additionally, we also calculated the average number of attacks on the honeybees’ appendages and bodies for each colony across the trials. We then performed Wilcoxon signed-rank paired tests to compare the average attacks on the bee’s appendages and body between the 2D and 3D colonies.

## Results

### Web architecture shapes collective hunting strategies

Our generalized linear mixed effects models show that the architecture of the capture web (2D vs. 3D) influenced the behaviour of *S. sarasinorum*. Spiders with 2D capture webs emerged faster from their nests in response to vibration (Mean ± SD, 2D colonies: 3.5 ± 2.23 *s*; 3D colonies: 4.98 ± 1.87 *s*) and attacked sooner than spiders possessing 3D webs (2D colonies: 11.70 ± 5.0 *s*; 3D colonies: 24.64 ± 9.31 *s*). Additionally, 2D colonies recruited significantly greater number of attacking spiders when compared to 3D colonies (2D colonies: 4.81 ± 1.14; 3D colonies: 3.08 ± 1.27; **Figure 2, Table 1**).

**Table 1:**
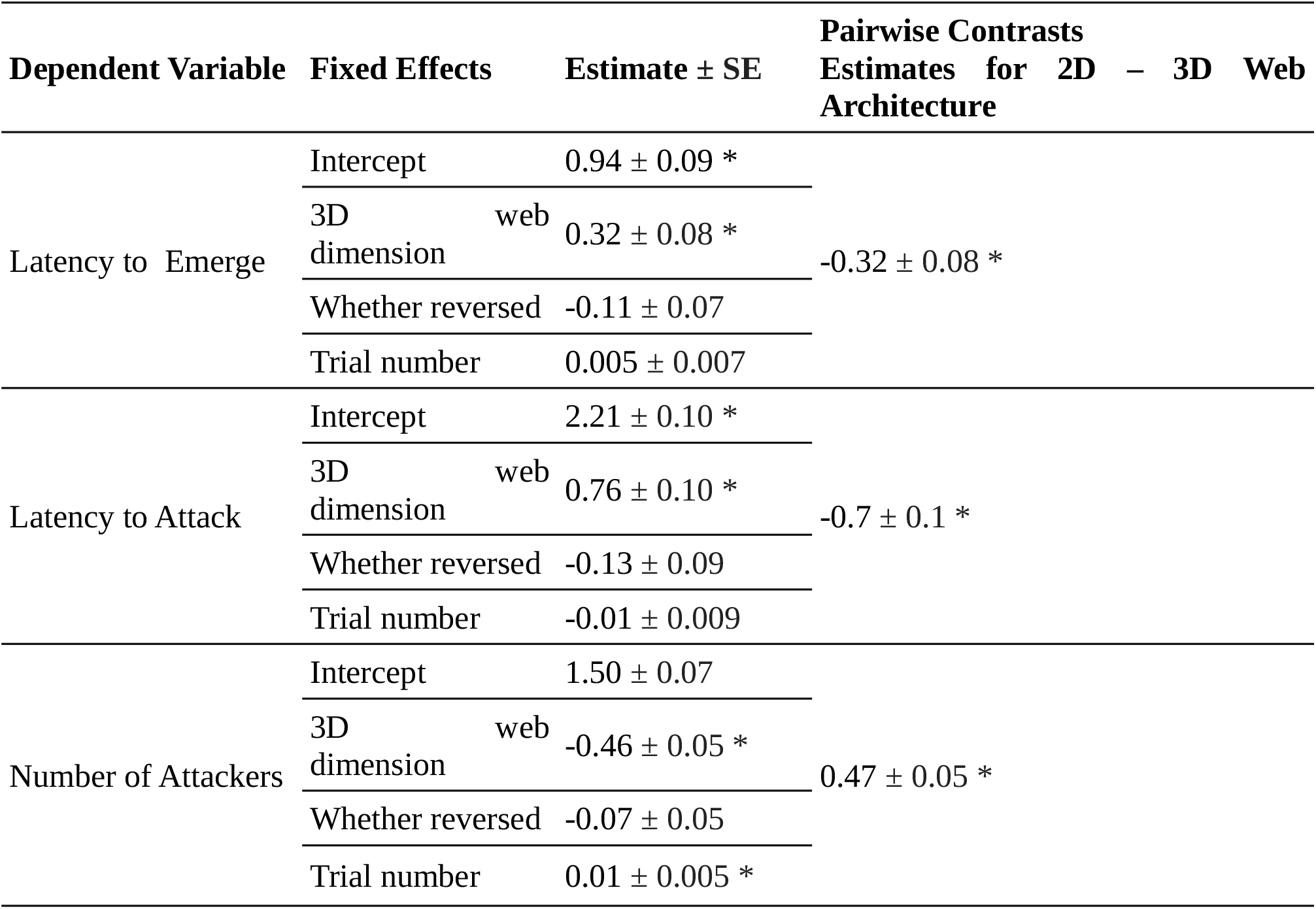
Estimates from three independent mixed models consisting of latency to emerge, latency to attack or number of attackers as the dependent variable. The estimates are in the (log)_10_ scale. * indicates significance at *P* < 0.05.

| Dependent Variable | Fixed Effects | Estimate ± SE | Pairwise Contrasts<br>Estimates for 2D – 3D Web<br>Architecture |
| --- | --- | --- | --- |
| Latency to Emerge | Intercept | $0.94 \pm 0.09$ * | $-0.32 \pm 0.08$ * |
| | 3D dimension web | $0.32 \pm 0.08$ * | |
| | Whether reversed | $-0.11 \pm 0.07$ | |
| | Trial number | $0.005 \pm 0.007$ | |
| Latency to Attack | Intercept | $2.21 \pm 0.10$ * | $-0.7 \pm 0.1$ * |
| | 3D dimension web | $0.76 \pm 0.10$ * | |
| | Whether reversed | $-0.13 \pm 0.09$ | |
| | Trial number | $-0.01 \pm 0.009$ | |
| Number of Attackers | Intercept | $1.50 \pm 0.07$ | $0.47 \pm 0.05$ * |
| | 3D dimension web | $-0.46 \pm 0.05$ * | |
| | Whether reversed | $-0.07 \pm 0.05$ | |
| | Trial number | $0.01 \pm 0.005$ * | |

**Figure 2:**
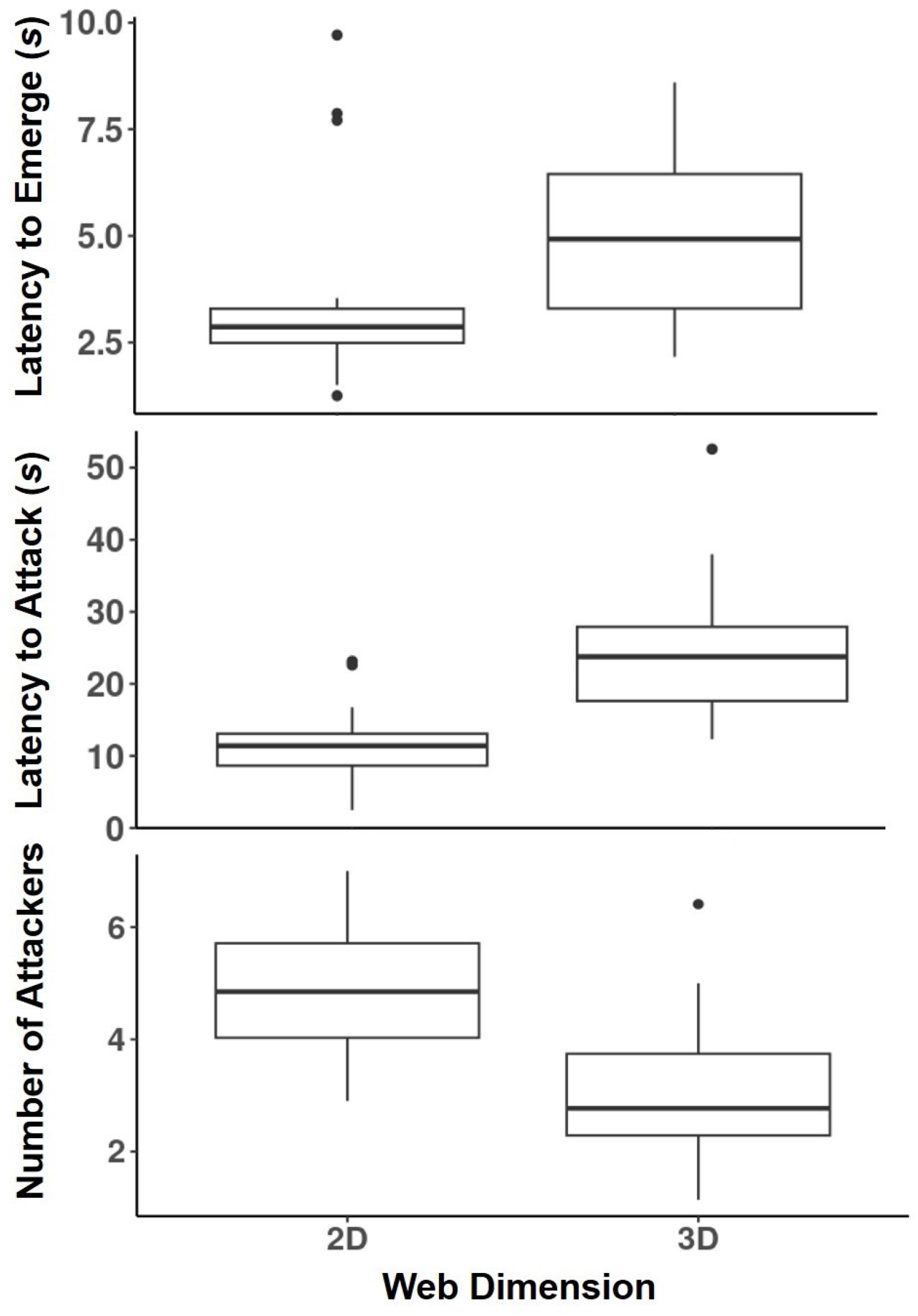
The figure illustrates the impact of web dimension (2D or 3D) on hunting behaviour in the spider species *S. sarasinorum*. Colonies with 2D capture webs exhibited significantly faster emergence and attack latencies, as well as a greater number of attackers, compared to colonies with 3D capture webs. The boxes depict the lower and upper quartiles, with whiskers representing data outside those quartiles, and filled circles indicating outliers. The internal horizontal lines within the boxes denote the median values.

### The 2D and 3D colonies employ distinct strategies to attack different body parts of their prey

In both 2D and 3D colonies, spiders predominantly attacked the appendages of honeybees (wings, legs, and antennae, 2D colonies: 82.90%; 3D colonies: 91.22%) than the body parts (head, thorax, abdomen, **Figure 3**). However, the percentage of spider attacks on the body increased after the bees were subdued (2D colonies: 68.1%; 3D colonies: 55.62%, **Figure 3**). This suggests that spiders approached dangerous prey like honeybees cautiously by avoiding their body parts. Interestingly, the attacks on the body parts of living bees were significantly higher for 2D colonies when compared to 3D colonies (2D colonies: 17.09%, mean of colony-wise averages across trials ± SD = 0.51 ± 0.25; 3D colonies: 8.77%, mean of colony-wise averages across trials ± SD = 0.27 ± 0.20, V = 18, P = 0.01, Wilcoxon signed-rank paired-samples test, **Figure 3**). This indicates that spiders on 2D webs were more prone to risk when attacking dangerous prey.

**Figure 3:**
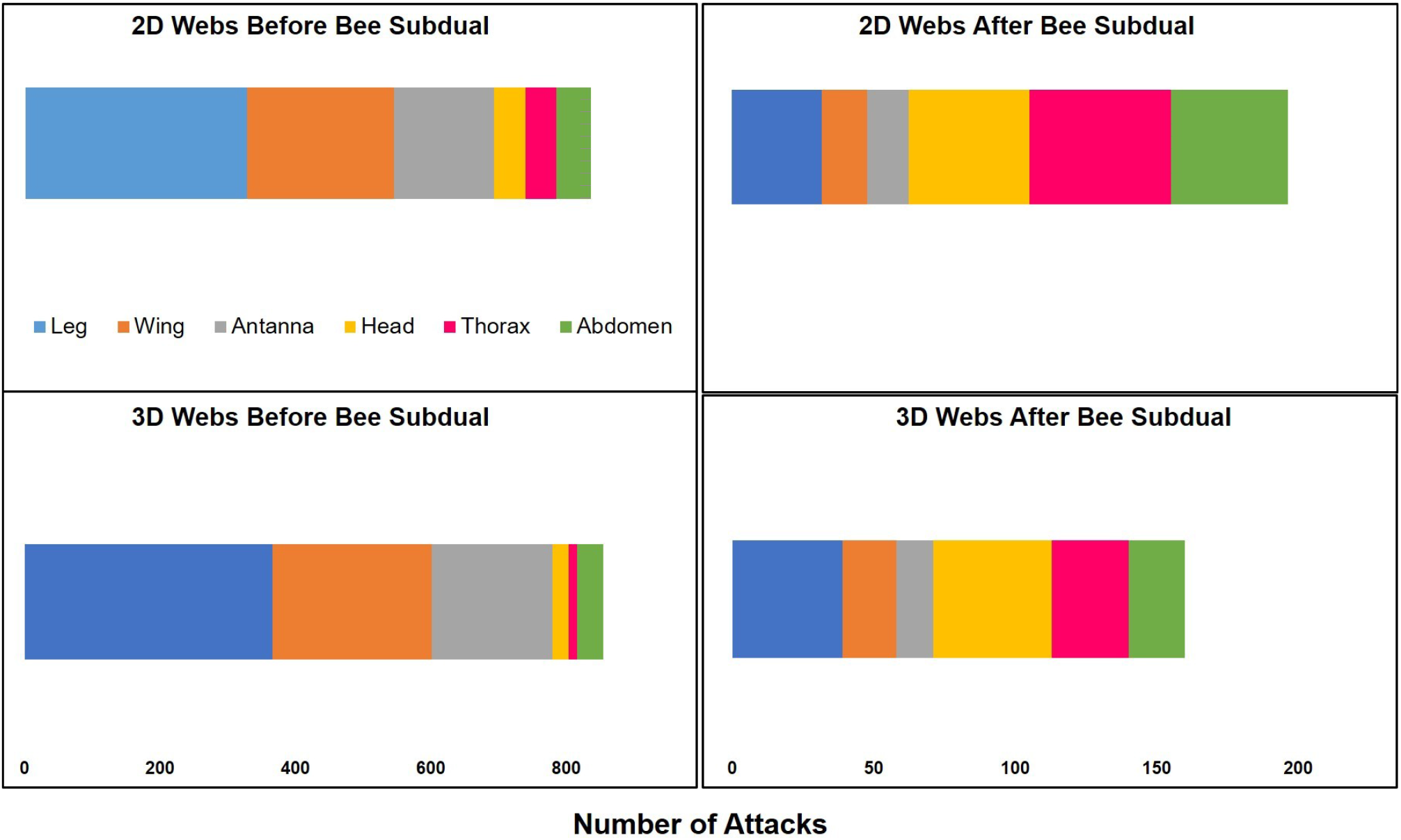
The figure shows the total number of attacks by *S. sarasinorum* spiders on various body parts of the honeybee (*Apis cerana*) before and after the bees were subdued. In colonies with 2D and 3D capture webs, the spiders attacked the bees’ appendages (legs, wings, antennae) more frequently than the body parts (head, thorax, abdomen) before the bees were subdued (left panels). However, the attacks on the body parts increased after the bees were subdued (right panels). Before the bees were subdued, the 2D colonies (top left panel) attacked the bees’ dangerous body parts more often than the 3D colonies (bottom left panel).

### Spiders hunting on two-dimensional webs more often fall victim to their prey

During our experimentation, we documented 7 spiders that succumbed to honeybee attacks across 6 different colonies. Of these, 5 out of 7 were documented in 4 separate 2D colonies, while the remaining 2 spiders belonged to 2 different 3D colonies.

## Discussion

We demonstrated that web architecture (2D vs. 3D web) significantly influences the collective behaviour of the social spider, *S. sarasinorum*. Spiders with 2D capture webs responded and attacked a simulated prey more quickly and in greater numbers compared to colonies with 3D webs. This shows that 2D webs can enhance the hunting success of social spider colonies, corroborating previous findings on the congeneric species *S. dumicola* (Kamath et. al 2019). Furthermore, we show that spiders with 2D capture webs were more likely to attack the dangerous body parts of the honeybee, *Apis cerana*, than those with 3D webs. This suggests that the web architecture can influence the risk-taking behaviour of *S. sarasinorum* when hunting potentially hazardous prey such as honeybees. Additionally, we also show that more spiders in 2D colonies succumbed to bee stings than those in 3D colonies. Collectively, our results show that human-induced habitat modification, which can alter web architecture, can in-turn impact the hunting behaviour and survival of *S. sarasinorum*. While 2D webs may confer benefits in hunting success, they also carry costs in terms of increased mortality. Further research is necessary to determine if the benefits of 2D webs outweigh the costs for these spiders

In our experiment involving living honeybees (experiment 1), only 7 spiders were stung to death by the bees. This low mortality could be attributed to the following two reasons: 1. we used only one prey species (*Apis cerana*) in our experiments, however, in the wild, mortality of spiders may be much higher due to their encounters with a variety of hazardous prey species. 2. Spiders approach dangerous prey like bees and wasps with caution and therefore, they avoid attacking their vital body parts (head, thorax, abdomen). A previous study (Parthasarathy & Somanathan 2019) supports our hypothesis by demonstrating that *S. sarasinorum* spiders avoided attacking the vital body parts of living honeybees, unlike their approach while hunting relatively harmless prey such as grasshoppers. However, the exact mechanism by which these spiders distinguish between dangerous and harmless prey remains unknown. One possible cue could be the nature of the vibratory signals on the web generated from the struggling prey. Dangerous prey like bees and wasps produce high-frequency vibrations while struggling on the web whereas more innocuous insects like crickets, grasshoppers, and butterflies generate lower-frequency, sporadic web vibrations. However, not all prey that emit high frequency vibrations on the web are dangerous to the spiders. For example, blow flies (family: Calliphoridae) are harmless yet they struggle in a way that resembles the high-frequency vibrations of bees and wasps. Interestingly, *S. sarasinorum* spiders handle blow flies with the same caution as they do with bees (Parthasarathy et al., unpublished), suggesting that *S. sarasinorum*, and possibly other spider species, treat any high-frequency web vibrations with care.

Despite their tendency to handle prey emitting high frequency vibrations with caution, spiders from 2D colonies attacked the bees’ vital body parts significantly more often than spiders from 3D colonies. This may explain the higher mortality due to bee attacks observed in the 2D colonies. The reason why spiders are less cautious while handling bees in 2D webs is unclear. One possibility is that the 2D webs, being simpler sheet-like structures in one plane, are less effective at ensnaring prey, such that bees’ appendages were not making enough contact with the web surface for the spiders to grasp. However, studies on other spider taxa show that 2D orb webs are better at ensnaring prey than 3D sheet webs (Rypstra 1982; Zschokke et al. 2006). Alternatively, the greater number of spiders attacking in 2D webs could direct some spiders to target the bee’s vital body parts, as the bee’s appendages are already occupied by other spiders. Finally, the *S. sarasinoru*m spiders, which typically build complex 3D capture webs in the wild, may not be fully adapted to hunting on the 2D webs constructed on man-made structures, given the relatively recent timescale of this habitat shift.

The reduced efficiency of 2D webs in ensnaring prey may explain the faster attack latencies and greater numbers of attacking spiders on these webs. Spiders on 2D webs need to act quickly to capture prey before it escapes, potentially requiring a collective effort from more individuals to subdue partially ensnared prey. Such trade-offs between attack speed and building webs that can retain prey longer are known in spiders (Zschokke et al. 2006). Alternatively, a more plausible hypothesis could be that the simpler structure of 2D webs, lying in a single plane, may allow for more efficient transmission of vibratory cues to the nests where spiders reside. The faster emergence time noted in 2D when compared to 3D colonies substantiates this. Furthermore, the relative simplicity of 2D webs may make it easier for spiders to navigate towards the prey, resulting in shorter attack latencies.

The vast majority of spider species construct 3D webs (Coddington & Levi 1991; Griswold et al. 1998), and one of the primary drivers behind the evolution of these 3D web forms from ancestral 2D webs may be the protection they offer against predators (Blackledge et al. 2003). Our study suggests an additional advantage of 3D capture webs: their ability to help spiders to safely handle risky prey, which could have contributed to the origin of this web form.

In conclusion, by experimentally manipulating the web architecture of lab-based spider colonies, we observed clear behavioural differences between 2D and 3D colonies. Notably, 2D colonies experienced higher mortality rates while hunting, suggesting that 3D capture webs provide spiders with better protection against dangerous prey. Further research is advocated to fully understand how web dimension impacts the survival, dispersal, and overall productivity of wild spider colonies.

## Acknowledgments

We thank Raghavendra Gadagkar for encouraging us to conduct this study and for providing the necessary facilities. We thank Rajbir Kaur, Shilpa Gowda and Anoushka Dasgupta for helping us with the experiments. This study was supported by the Centre for Excellence in Mathematical Biology, Sri Sathya Sai Institute of Higher Learning, Puttaparthi, India. BP was supported by the National Postdoctoral Fellowship from Science and Engineering Research Board (PDF/2019/001096).

## Author Contributions

BP: Conception, experimental design, supervision, performing experiments, data acquisition, formal analyses and writing the manuscript. NYS: Performing experiments, data acquisition, data curation and editing the manuscript. VSA, CVS and MVSK: Performing experiments, data acquisition and data curation. KKVD: Supervision and fund acquisition. All authors read and approved the final version of the manuscript.

## Supplementary Section

**Supplementary Table 1:**
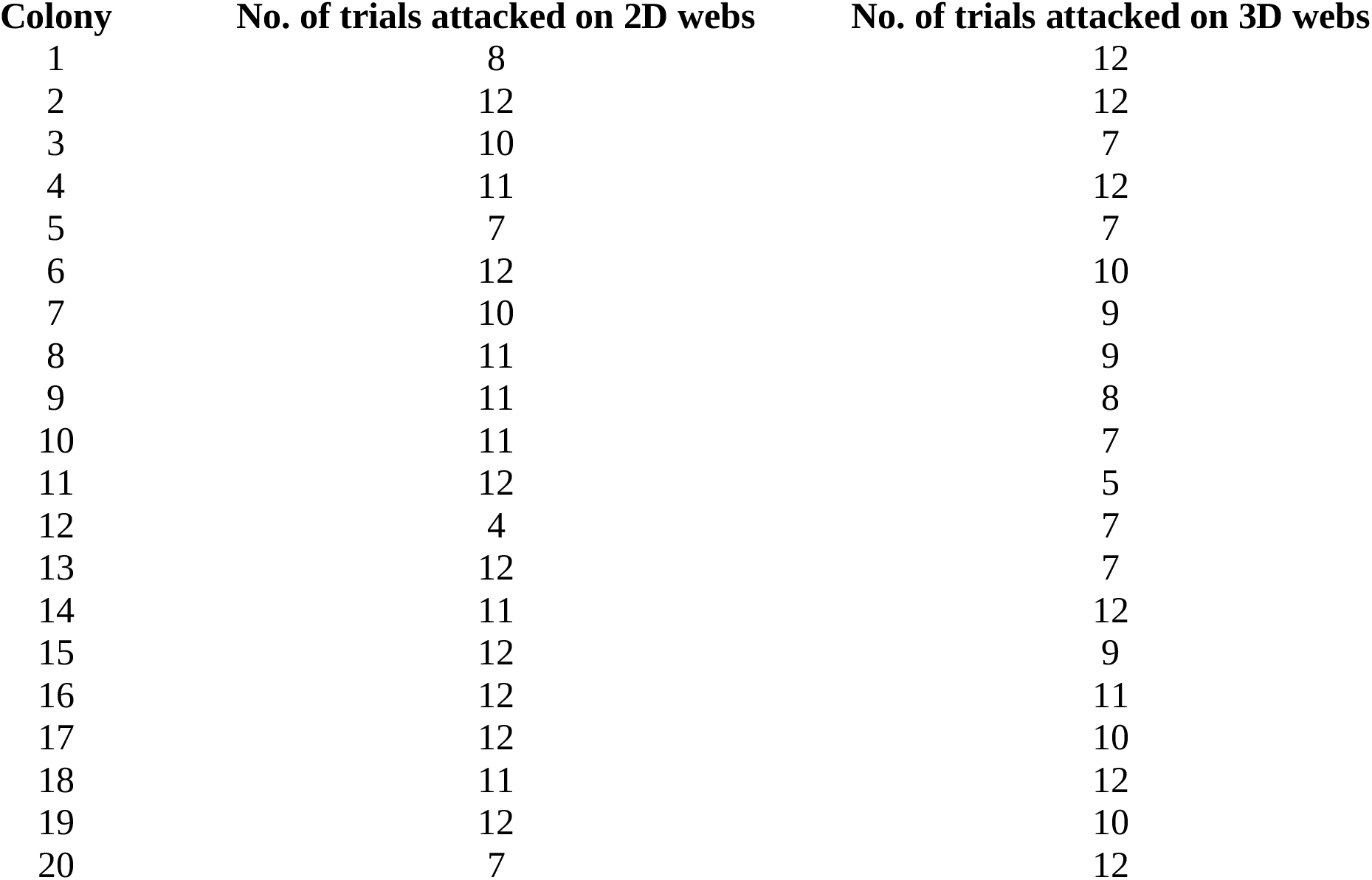
The table presents the number of trials in Experiment 2 where each spider colony attacked a simulated prey on 2D and 3D capture webs. A total of 12 trials were conducted for each colony. Trials were abandoned if a colony failed to respond to the vibration of the simulated prey within 2 minutes.

