## Supplementary Table 1 for "Extended Phenotype Influences Collective Behaviour and Survival in a Social Spider"

### Supplementary Section

**Supplementary Table 1:** The table presents the number of trials in Experiment 2 where each spider colony attacked a simulated prey on 2D and 3D capture webs. A total of 12 trials were conducted for each colony. Trials were abandoned if a colony failed to respond to the vibration of the simulated prey within 2 minutes.

| Colony | No. of trials attacked on 2D webs | No. of trials attacked on 3D webs |
|---|---|---|
| 1 | 8 | 12 |
| 2 | 12 | 12 |
| 3 | 10 | 7 |
| 4 | 11 | 12 |
| 5 | 7 | 7 |
| 6 | 12 | 10 |
| 7 | 10 | 9 |
| 8 | 11 | 9 |
| 9 | 11 | 8 |
| 10 | 11 | 7 |
| 11 | 12 | 5 |
| 12 | 4 | 7 |
| 13 | 12 | 7 |
| 14 | 11 | 12 |
| 15 | 12 | 9 |
| 16 | 12 | 11 |
| 17 | 12 | 10 |
| 18 | 11 | 12 |
| 19 | 12 | 10 |
| 20 | 7 | 12 |
